# Navigating *in vitro* bioactivity data: investigating available resources using model compounds

**DOI:** 10.1101/248898

**Authors:** Sten Ilmjärv, Fiona Augsburger, Jerven Tjalling Bolleman, Robin Liechti, Alan James Bridge, Jenny Sandström, Vincent Jaquet, Ioannis Xenarios, Karl-Heinz Krause

## Abstract

Modern medicine and an increasingly complex environment contribute to exposure of humans to a large number of chemical compounds, that can potentially be toxic. Although widely used, compound testing in animals has important limitations. *In vitro* testing provides a promising alternative. However, because of the relative inaccessibility and fragmentation of available data, the *in vitro* approach largely underperforms its potential. The aim of this study is to investigate how available public online resources (tools and databases) support accessing and distribution of *in vitro* compound data. We examined 19 public online resources, mapped their features, and evaluated their usability with a set of four model compounds (*aspirin, rosiglitazone, valproic acid*, and *tamoxifen*). By investigating compound names and identifiers, we observed extensive variation and inconsistencies in available resources: the synonyms were different, compounds’ structural identifiers (InChI, InChIKey, SMILES and IUPAC systematic name) underperformed in omics databases, identification of compound related metadata (e.g. concentrations used in the experiments) from omics experiments was complex and none of the available resources clearly distinguished between *in vivo* and *in vitro* data. In addition, we estimated accessibility of selected public resources using computational queries. Only a few public resources provided access to compound-related data using semantic web technology. The general quality of experiment annotations created further difficulties in identifying data of interest. Therefore, we identified several standardized ontologies with potential to provide an increased accuracy for extensive data retrieval of *in vitro* compound data. Furthermore, using the examples of our model compounds, we provide recommendations on the use of ontologies by suggesting specific ontology terms to annotate *in vitro* experimental data when being published.

## Introduction

The number of chemical compounds in public databases is growing and experimental data concerning these compounds are accumulating. As of today, both the CAS Registry^SM^ and PubChem^1^ contain more than 90 million compounds with many new compounds being added each day. Among these compounds are drugs, chemicals, environmental contaminants and toxins, all of which potentially could elicit effects that could have implications on human health and/or environment. Most of these compounds have not been characterized from a toxicological point of view. Indeed, the US based ToxNet database Hazardous Substances Data Bank (HSDB)^2^ where an expert panel reviews individual compounds, contains less than 6000 records (as of 12.12.2017). Alternatively, ChEMBL^3^, which predominantly consists of literature extracted bioactivity data, contains around 1.73 million distinct compounds (as of 12.12.2017). The difference in the number of compounds registered and the number of compounds whose toxic effect is well characterized, demonstrates the limited capacity of current methods to assess a compound’s bioactivity in a living system. *In vitro* test systems with high-throughput performance and potential scalability aim to bridge that gap. It will further increase the amount of data and provide new knowledge of compound’s biological properties.

Animal testing of compound toxicity and efficacy is requested for drug development, but it is slow, expensive and raise concerns about animal welfare^4^. Alternatives to animal experiments is strongly encouraged according to the 3R (replacement, reduction and refinement) principles. Therefore, large-scale initiatives have been launched to investigate compound’s toxicity in rapid, quantitative and scalable *in vitro* systems (e.g. Tox21^5^ and EU-ToxRisk^6^). These bioassays can measure endocrine disruption^7^, the generation of reactive oxygen species (ROS)^8^ or changes in gene expression of biomarkers in developing and mature neurons^9^. These assays represent different aspects of the human biology where an aberration of homeostasis can suggest adverse consequences to the human health. Compared to animal models, *in vitro* bioassays allow studying concentration-response relationships over a large concentration range, including those representative of human exposure^4^. By combining compound’s structural information, known molecular properties and data from bioassays and omics experiments, researchers can better describe compound effect pathways for a systematic understanding.

Presently there is no simple way to access *in vitro* compound data in a quick and synoptic manner. Instead, data is fragmented across many different resources^10^ and interested parties need to invest invaluable time and effort to develop an expertise in order to navigate these systems efficiently^11^. The diversity of compound synonyms and its identifiers, lack of precise metadata and annotations, can lead to false conclusions and to difficulties identifying the compound correctly after it has been published^12^. To improve the reproducibility of experimental results and to test new hypotheses (e.g. development of predictive computational models), availability and accessibility of raw data is crucial.

In this study, we analyzed 19 public resources that potentially allow a researcher to investigate compound-specific *in vitro* data. We demonstrate that modest adoption of semantic web technologies and poor annotations of experimental metadata, in particular when describing experimental conditions, represent a major obstacle for high quality data integration and its reusability. Lastly, we provide insight how standardized ontologies could improve the annotation of compound related experimental data.

## Materials and methods

### Choice of model compounds for resource analysis

In order to search the resources and to identify their potentials and pitfalls, we chose four model compounds that were used to evaluate public resources throughout this study (Table 1): *rosiglitazone*, *valproic acid*, *aspirin* (*acetylsalicylic acid*), and *tamoxifen*. Below the motives to why these were chosen as model compounds are listed:

**Table 1.**
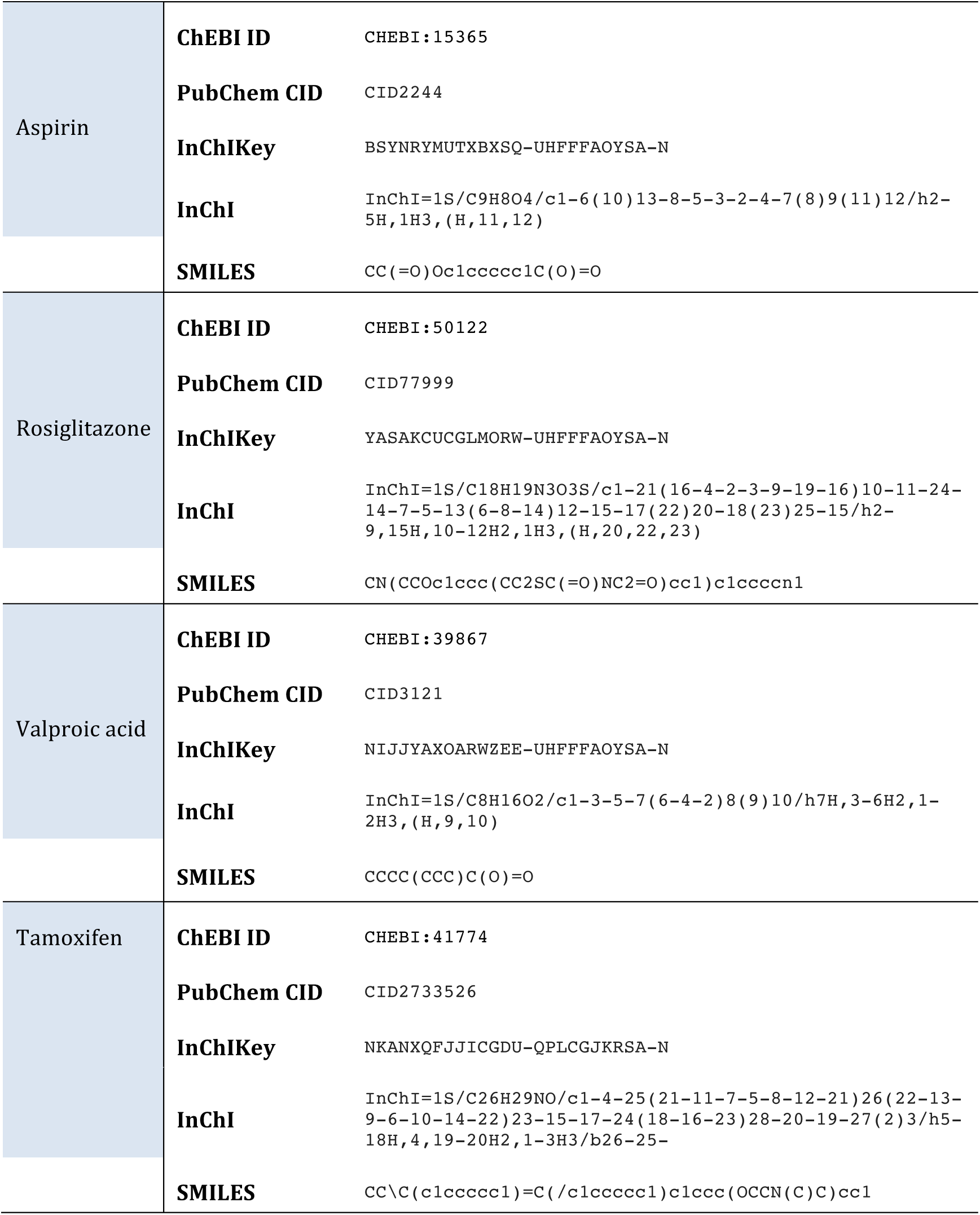
Table of model compounds used in the study and their identifiers.

*Tamoxifen* is a commonly used drug to treat hormone-dependent breast cancers. *Tamoxifen* was included in the study because of the several known side-effectsl^13^ and its relatively complex structure.

*Valproic acid* is an antiepileptic drug which has been in clinical use for about half a century. Its mechanisms of action are multiple and include inhibition of histone deacetylases^14^. Its therapeutic use is well accepted but it has been found to cause severe developmental toxicity if given to pregnant women^15^. The history of the drug made it interesting for us to study if later discoveries of toxic effects were translated to better data representation and to additional experiments performed with the compound.

*Aspirin* (*acetylsalicylic acid*) is one of the oldest synthetic drugs. Its mechanism of action, inhibition of prostaglandin synthesis through acetylation of platelet cyclooxygenase (COX)^16^, makes it an anti-inflammatory, analgesic and antipyretic drug. Aspirin was interesting for our study since for a long time it has been widely used as a readily available painkiller, thus potentially having many historical synonyms and experimental data. Of note, the name aspirin itself derives from a trade name.

*Rosiglitazone* is an antidiabetic drug of Peroxisome Proliferator-Activated Receptors (PPAR) agonists family, introduced into the market in 1999. It was included in the analysis because compared to other model compounds it is relatively new. Also, similar to *tamoxifen*, the complex chemical structure of *rosiglitazone* enabled us to study the structural identifiers of more intricate molecule.

### Overview of databases containing compound information

To get an overview of public resources containing compound information, both compound specific resources (e.g. databases containing physicochemical properties such as structure) and omics databases were investigated using our model compounds. Primarily, we focused on resources that could potentially accommodate or link to *in vitro* compound data. Downstream information such as target interactions and pharmacological inference were also considered as they can be derived from *in vitro* experiments and hence raw data could potentially be backtracked. Resource features included in our analysis involved the scope of supported compound identifiers and search functionalities, accessibility of raw data, support for ontologies and programmatic data access.

The public resources we used in our study were PubChem^1^, ChEMBL^3^, ChEBI^17^, Chemistry Dashboard (CompTox)^18^, ChemSpider^19^, BioSamples^20^, ArrayExpress^21^, ExpressionAtlas^22^, PRIDE PRoteomics IDEntifications (PRIDE)^23^, Human metabolome database (HMDB)^24^, The Toxin and Toxin Target Database (T3DB)^25^, Gene Expression Omnibus (GEO)^26^, UniProt^27^, BindingDB^28^, DrugBank^29^, ZINC^30^ and three of the Toxicology Data Network (TOXNET) databases: Hazardous Substance Data Bank (HSDB)^2^, ChemIDPlus^31^, and Comparative Toxicogenomics Database (CTD)^32^ (Table 2).

**Table 2.** A list of resources used in the study, their categorization and the number of estimated compounds in these resources at the time of the study.

| Database | No. of compounds | Checked at | Data type |
| --- | --- | --- | --- |
| ArrayExpress | - | - | Raw |
| BindingDB | 635 301 | December-2017 | Curated |
| BioSamples | - | - | Raw |
| ChEBI | 53 495 | December-2017 | Curated |
| ChEMBL | 1 735 442 | December-2017 | Curated |
| ChemIDPlus | 418 376 | December-2017 | Curated |
| ChemSpider | ~ 62 million | December-2017 | Curated |
| CompTox | ~ 760 000 | December-2017 | Raw/ Curated |
| CTD | 15 381 | December-2017 | Curated |
| DrugBank | 10 562 | December-2017 | Curated |
| ExpressionAtlas | - | - | Raw/Curated |
| GEO | - | - | Raw |
| HMDB | 114 100 | December-2017 | Curated |
| HSDB | 5929 | December-2017 | Curated |
| PRIDE | - | - | Raw |
| PubChem | > 94 500 000 | December-2017 | Raw/Curated |
| T3DB | 3 673 | December-2017 | Curated |
| UniProt | - | - | Curated |
| ZINC15 | > 100 000 000 | December-2017 | Curated |

### Model compounds, compound naming and identifiers

To demonstrate the nomenclature and data retrieval complexity of public resources, we used ChEBI as our reference database since it is extensively curated and ChEBI ontology^17^ was used in several other resources (eg. PubChem and ChEMBL). In addition, ChEBI contained an ontology term for all of our four model compounds (Table 1). For an overview of compound names, synonyms and identifiers, we examined the public resources in Table 2. Our scope of identifiers and registry numbers included Simplified Molecular Input Line Entry System (SMILES), International Chemical Identifier (InChI), International Chemical Identifier Key (InChIKey) and International Union of Pure and Applied Chemistry (IUPAC) name. SMiles ARbitrary Target Specification (SMARTS) were not included in our analysis since they are not shown in ChEBI. Also, we excluded Chemical Abstract Service Registry Number (CASRN) because the accuracy of CASRN in the public domain are not absolute and reliable information can only be accessed by paid services provided by Chemical Abstract Service (CAS)^10^.

### Data access and integrative initiatives

The ability of a resource to integrate data and compound information was evaluated by searching cross-references between resources using the selected model compounds. To assess data sharing and access to public data collections of the selected resources, we studied the use of representational state transfer (REST or RESTful) application programming interface (API) and resource description framework (RDF) technologies.

### Analysis of ontology usage

To support the use of data cross-referencing from individual experiments we evaluated if and to what extent, the public resources utilized ontologies in their data annotation and presentation. With the exception of Cellosaurus^33^, the rest of the ontologies we considered important were included either in the OBO Foundry^34^, NCBI BioPortal^35^ or Ontology Lookup Service by EBI^36^. Although there might be other ontologies not included in these collections, they were considered less visible and therefore less likely to be taken up by the research community.

## Results

### Description and interconnectivity of public resources containing compound data

Here, our aim was to study what are the public resources containing compound information and if one resource could be used to identify a compound in another resource. Broadly, compound related information can be separated into two categories: 1) knowledge that has been manually or automatically curated from publications, which often result from published or unpublished raw data (e.g. half maximal effect concentration (IC_50_) values or compound interactions with targets), and 2) the raw data itself, either from omics experiments that interrogate many potential interactions simultaneously or from dedicated bioassays that measure a specific endpoint over a range of concentrations. Therefore, the resources can be classified as either containing curated information from scientific publications or some other reliable documents (examples include Hazardous Substance Data Bank (HSDB), ChEMBL), or databases that contain raw numerical data dedicated to certain data types (examples include ArrayExpress, PRIDE). In the case of PubChem, it supports both curated information and raw data of bioassays as uploaded by its users. An overview of databases with the type of information they contain can be found in Table 2.

We observed, that often a resource is actually reusing data from other databases, thus combining information from several resources, or providing cross-references to them. As such, DrugBank, PubChem, and ChEBI had the highest number of incoming connections from other resources and PubChem, ChEMBL, ChEBI and ChemSpider had the highest number of outgoing connections, connecting roughly to half of the public resources in our study list (Figure 1). Most of the resources had cross-references to another resource, especially when they contained data curated from scientific documents. However, with the exceptions of UniProt, ChEBI and ChEMBL, no compound-specific resources had cross-references to databases containing data from omics experiments. At the time of the study we were able to identify references in ChEBI to *in vitro* raw data in ArrayExpress for three out of four model compounds (*rosiglitazone*: E-GEOD-5509, E-GEOD-5679, E-GEOD-30147; *valproic acid*: E-GEOD-1615, E-GEOD-14973, E-GEOD-23909, E-TABM-903, E-TABM-1205, *tamoxifen*: E-GEOD-2225, E-GEOD-12665, E-MTAB-5319, E-TABM-562). For *aspirin*, we were able to find 73 datasets in ArrayExpress, representing both *in vivo* and *in vitro* experiments that have until now not been cross-referenced in ChEBI.

**Figure 1.**
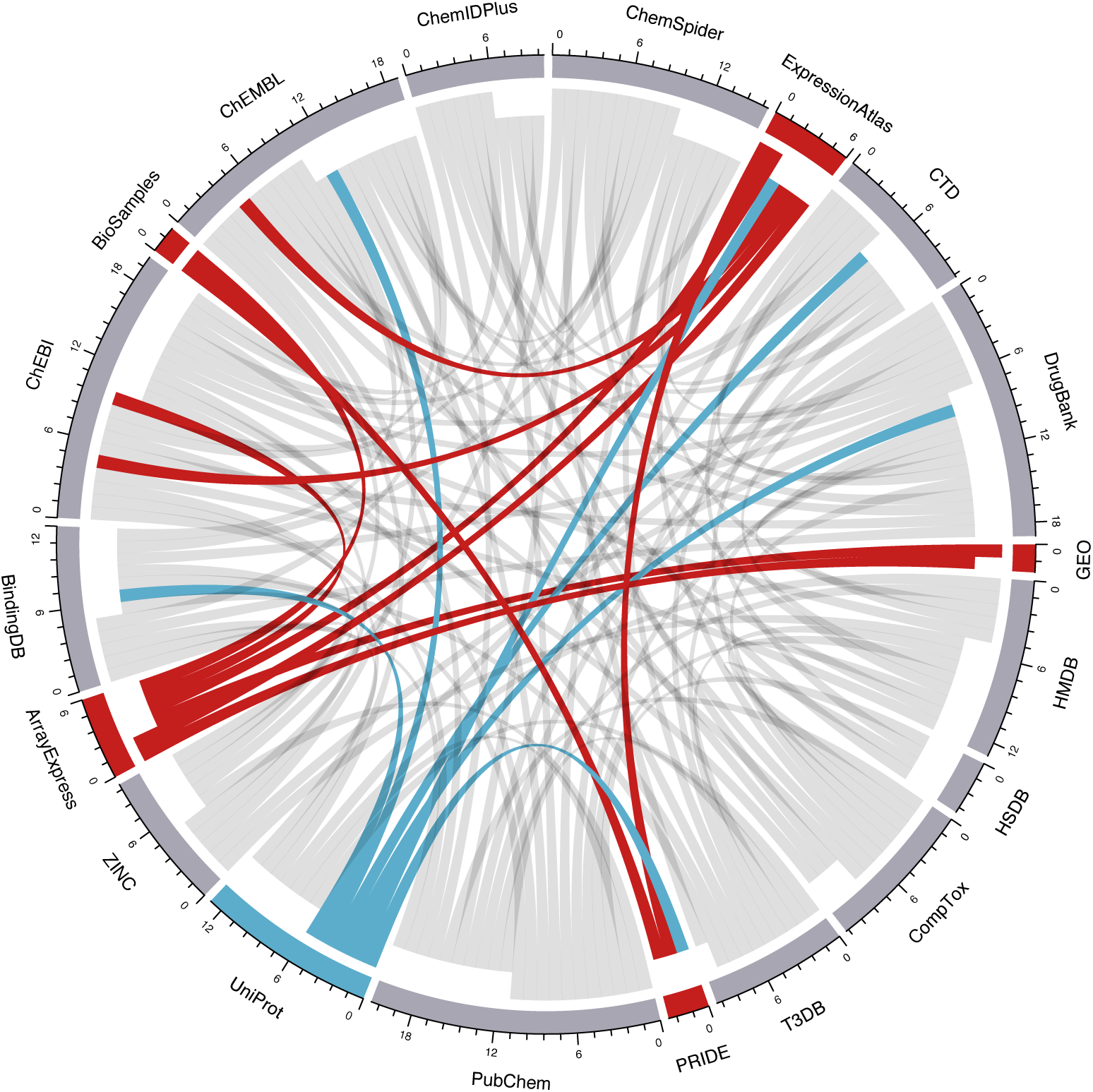
Representation of cross-references (representing either direct links to other resource or sampled data from the other resource) between resources in a chord diagram. Grey resources are depicting compound-specific resources whereas, the red ones depict omics resources. UniProt is neither compound-specific or omics resource and is therefore colored blue. Outgoing connections are closer to the circumference whereas incoming connections are further from the circumference. The numbers on the scale represent the combined number of both incoming and outgoing connections for every resource.

### Identification of data in compound-specific public resources

Here, our aim was to study different ways a chemical compound is identified between various public resources. Indeed, a chemical compound can be associated with many identifiers, such as a trade name, a generic name, a registry number, a unique database identifier (e.g. PubChem CID or ChEMBL ID) and its structure-derived representations, referred here as structural identifiers (SMILES, InChI, InChIKey – different types of compound identifiers are shown in Table 3). Potentially any of these can be used to search a compound from an online public resource. However, we observed that not all compound synonyms and identifiers are identical in different resources. As an example, the compound *rosiglitazone* contains 151 depositor-supplied synonyms in the PubChem^37^, whereas in ChEBI only two synonyms were provided^38^. Consequently, the PubChem depositor-supplied synonym for *rosiglitazone* termed *Gaudil* failed to recognize the compound in ChEBI.

**Table 3.** A table of compound identifiers, their features and examples. A better overview of compound identifiers is provided by Warr, 2011^39^. Of note, there are several structure-based identifiers and not all of them allow for substructure searches. In addition, some structure-based identifiers (e.g. InChIKeys) are better suited for cross-referencing purposes.

| Identifier | Standardized | Unique representation of a compound | Machine-readable structure notation | Often used in annotating data | Permits substructure search | Can be used for cross-referencing | Examples |
| --- | --- | --- | --- | --- | --- | --- | --- |
| Compound name |  |  |  | ✓ |  |  | Rosiglitazone |
| Trade name |  |  |  | ✓ |  |  | Avandia |
| Compound database identifier |  | ✓ |  |  |  |  | CHEMBL121 |
| Registry number |  | ✓ |  | ✓ |  |  | 122320-73-4 (CAS number) |
| Chemical structure drawing | ✓ | ✓ |  |  | ✓ |  |  |
| Structure-based identifier | ✓ | ✓ | ✓ |  | ✓ | ✓ | SMILES, InChI, InChIKey, IUPAC name |

To further illustrate the complications in searching information for a compound from several distinct databases, we first used compound names of four model compounds to investigate compound specific resources (BindingDB, ChEBI, ChEMBL, ChemIDPlus, ChemSpider, CompTox, CTD, DrugBank, HMDB, HSDB, PubChem, T3DB and ZINC15) using free-text based search. We were able to find records for all four model compounds in all of the resources (with one exception of *rosiglitazone* that was not found in T3DB) and we recorded their structural identifiers where available (Supplementary Table 1-4). Interestingly, we observed that not all structural identifiers were identical between the resources. That was especially true for SMILES where from the eleven resources that reported SMILES, we found 8 different SMILES for *rosiglitazone* and *tamoxifen*, 5 for *aspirin* and 3 for *valproic acid* (Supplementary Table 1-4). The results were better for InChIKeys (established by International Union of Pure and Applied Chemistry (IUPAC)), where we observed a single identifier for both *aspirin, valproic acid* and *tamoxifen* but three different InChIKeys for *rosiglitazone*. Things were more complicated with IUPAC systematic names since in several resources it was unclear if the reported name was actually a IUPAC systematic name and therefore these resources were discarded from the analysis. Consequently, IUPAC name was recorded only from ChEBI, ChemSpider, CompTox, DrugBank, HMDB, PubChem and T3DB. Even though few resources reported IUPAC systematic names, we observed large variability among them. More specifically, we found 3 different names for *aspirin,* 4 for *rosiglitazone,* 1 for *valproic acid* and 5 for *tamoxifen* (Supplementary Table 1-4).

These results highlight that InChIKeys among various databases are more unique compared to SMILES or IUPAC systematic names. To that end, a global resource UniChem has provided a cross-referencing service that connects 38 individual database identifiers of various resources using InChIKeys^40^. However, this service is only useful when one already knows the compound’s database identifier or the InChIKey. Currently, it cannot be used with other structural identifiers or with compound names.

### Identification of compound data from omics databases

The identity of the chemical compounds can be ambiguous, since compounds are often mentioned by name without the accompanying structure representations^12^. This is especially true for omics resources where data uploaded by the researchers can be annotated with different synonyms of the same compound. To investigate this issue, we utilized a web-based free-text search to research omics data from ArrayExpress, ExpressionAtlas, BioSamples, GEO and PRIDE resources using compounds’ structural identifiers (SMILES, InChI and InChIKey) as reported in ChEBI. From all the resources, we were able to retrieve data at least for two model compounds using compound names (Supplementary Table 5). In addition, the IUPAC systematic names of *aspirin, rosiglitazone* and *valproic acid* retrieved datasets from ArrayExpress, BioSamples and GEO. Interestingly, in BioSamples, we were able to retrieve datasets for *valproic acid* also with SMILES. However, these datasets actually corresponded to the sodium salt of *valproic acid,* that has a slightly different SMILES representation in ChEBI compared to *valproic acid*. Moreover, these samples were not retrieved when the compound name was used instead.

The latter highlights that the best way to identify compound related data from omics resources is to use compound names, which would require the researcher to consider all of the compound synonyms. Therefore, to estimate the variability of compound annotation in sample labels, we retrieved the name, synonyms and structural identifiers for each of the four model compounds from the ChEMBL public SPARQL endpoint (the code is available at https://github.com/whetlake/ivcdp). These were then used to identify samples and sample labels in the BioSamples database using BioSamples public SPARQL endpoint. For *rosiglitazone* and *tamoxifen* only the samples with the respective name was found in any of the sample labels. For aspirin, samples were found using both *aspirin, asparin, asprin, levius* and *measurin* (Supplementary Table 6-9). The term *measurin* can also be derived from sample labels that contain the word *measuring,* making it highly susceptible to retrieving false samples, not associated with aspirin. Surprisingly, the compound name *acetylsalicylic acid* highlighted in ChEBI was not found in any of the sample labels. *Valproic acid* retrieved results also for *valproate, depakote* and *44089*. The latter is a synonym of *valproic acid* in ChEMBL but none of the associated samples were actually associated to *valproic acid*. Similar to web-based free-text search, in our analysis, no samples contained structural identifiers of the model compounds within the sample labels. Also, all the samples retrieved were unique i.e. alternative compound labels were not used to annotate the same sample. In addition, care must be taken when using free-text based approach to programmatically search for compound data. Indeed, we were able to retrieve samples where sample labels indicated a negation of the compound, such as “*No rosiglitazone present*” (SAMEA862998^41^).

### Identification of *in vitro* compound data from public resources

With the exception of ZINC15, none of the public resources clearly distinguished compounds whose effect has been measured in *in vitro* experiments. In contrast, ZINC15 reports a subsets of compounds that have been reported or inferred active in *in vitro* direct binding assays^42^. Another approach to identify *in vitro* data in public resources is to browse the study description for references of *in vitro* experiment related keywords like “*in vitro*”, “cell-line” or specific cell-line names (e.g. “HeLa”). For example, PubChem, ChEMBL, ChEBI, ArrayExpress, ExpressionAtlas, GEO, BioSamples and PRIDE can be used to map these keywords with sample descriptions using free-text search. As such, we were able to retrieve a bioassay record from PubChem with the title “*In Vitro* Cytotoxicity Against Ovarian Carcinoma Cell Line CH1”^43^ which clearly represents an *in vitro* experiment. ChEMBL provides a more precise search, which allows to retrieve data on compounds associated with specific cell-lines or *in vitro* assays.

In addition, we can identify *in vitro* data by using concentration units associated with molarity or molality. For example, we used previously identified microarray samples from ArrayExpress experiment E-GEOD-5679 where dendritic cells were treated with *rosiglitazone*^44^. From the full experiment description, we identified that *rosiglitazone* was used at 2.5 μM concentration level, indicating that this was indeed an *in vitro* experiment. However, samples themselves had no attribution of concentration unit in neither ArrayExpress^44^ or BioSamples (SAMEG19226^45^). Since the actual concentration could only be retrieved from the experiment description or from the original publication, this approach is currently not scalable to large-scale automatic identification of *in vitro* data, thus limiting its use.

In summary, this analysis demonstrates that presently no concrete way to specifically identify *in vitro* data using free-text based searches exists in any of the studied resources. Although ChEMBL provides identification of compound information related to cell-lines and *in vitro* assays, one would have to cover all available cell-lines and assays in order to get a complete overview of the available *in vitro* data.

### Accessing *in vitro* compound data from public resources programmatically

To reuse already published data, relevant data need to be identified correctly and made available to the researcher. Therefore, most public resources provide access to data through a bulk download (Table 4), which can be very large (e.g. the PubChem compound database in compressed ASN format is about 60GB, as of March 2017). Alternatively, several public resources also provide online programmatic data access. Here, we researched the utilization of RESTful API (Representational state transfer application programming interface) and Resource Description Framework (RDF) technologies by the selected public resources.

**Table 4.** A list of resources used in the study and their support for different data access.

|  | Bulk download | RESTful API | RDF support | SPARQL endpoint | Substructure and/or similarity search |
| --- | --- | --- | --- | --- | --- |
| ArrayExpress | ✓ | ✓ |  |  |  |
| BindingDB | ✓ | ✓ |  |  | ✓ |
| BioSamples |  | ✓ | ✓ | ✓ |  |
| ChEBI | ✓ |  | ✓ |  | ✓ |
| ChEMBL | ✓ | ✓ | ✓ | ✓ | ✓ |
| ChemIDPlus | ✓ | ✓ |  |  | ✓ |
| ChemSpider |  | ✓ | ✓ |  | ✓ |
| CompTox | ✓ | ✓ |  |  |  |
| CTD | ✓ |  |  |  |  |
| DrugBank | ✓ | ✓ |  |  |  |
| ExpressionAtlas | ✓ | ✓ | ✓ | ✓ |  |
| GEO | ✓ |  |  |  |  |
| HMDB | ✓ |  |  |  | ✓ |
| HSDB | ✓ | ✓ |  |  |  |
| PRIDE | ✓ | ✓ |  |  |  |
| PubChem | ✓ | ✓ | ✓ |  | ✓ |
| T3DB |  |  |  |  | ✓ |
| UniProt | ✓ | ✓ | ✓ | ✓ |  |
| ZINC15 | ✓ |  |  |  | ✓ |

In a RESTful query, the data request is constructed into a single URL. This makes RESTful API simple to use and platform independent. Out of the 19 resources in our study, 10 provided free access to their RESTful API (Table 4), whereas access to DrugBank API was provided for a fee. Contrary to RESTful API where no data is stored locally, the data in RDF compatible format can be used for bulk download and subsequent introduction into a local database for query execution. However, a preferred way of data acquisition is through a public SPARQL endpoint, which facilitates querying the service provider directly, thus always retrieving the most up-to-date data. Here we observed that only BioSamples, ChEMBL, ExpressionAtlas and UniProt provide a public SPARQL endpoint (Table 4). Acquiring data using a SPARQL endpoint can be slower compared to RESTful data access, since the latter is better optimized for specific, recurrent query requests. However, SPARQL queries have the benefit of being highly customizable by the researcher and therefore they provide the flexibility that will cater to the researchers’ unique needs. Also, since RDF is an inherent part of the “linked data” concept, it can be used to find relationships between datasets in different resources and is therefore extremely useful for data integration purposes, such as connecting compounds effect in one resource to its physicochemical properties in another.

### Identification of relevant ontologies to annotate *in vitro* compound data

Ontology is a collection of specifically defined and controlled vocabulary consisting of ontology terms. Ontologies allow to formally describe concepts and to define the relationship between them^46^. Individual ontologies can consist from few to many thousands of ontology terms. For example, ChEBI ontology is made up over 100 000 ontology terms. By using ontologies, the researcher can create better targeted and more precise queries within the public resources. This necessitates that the researcher is able to identify and apply relevant ontology terms for their data. For this purpose, we identified three resources that allow to browse and search for specific ontologies and ontology terms: i) OntoBee^47^ is the default service containing a collection of ontologies as provided by the Open Biological Ontology (OBO) foundry^34^, as well as ii) BioPortal^35^ and iii) Ontology Lookup Service (OLS)^36^ which are ontology browsers developed by NCBI and EMBL-EBI, respectively. These services can be used to browse through a number of different ontologies, to view relations between terms and retrieve definitions for biomedical vocabulary. Currently (as of 20.12.2017), OLS contains 204, OntoBee 193 and BioPortal 678 ontologies. Important to our study, we identified 10 ontologies that are useful in relation to chemical compounds and can be used for annotating as well as programmatically accessing *in vitro* compound data (Table 5). As such, CHEMical INFormation ontology (CHEMINF)^48^ provides ontology terms for resource and structure identifiers. For example, InChIKey can be identified with a specific Universal Resource Identifier (URI) (http://semanticscience.org/resource/CHEMINF_000059) in CHEMINF ontology.

**Table 5.** Overview of ontologies that can be used to annotated or identify in vitro data.

|  |  |
| --- | --- |
| **ChEBI** | Chemical Entities of Biological Interest<sup>17</sup><br>Highly curated controlled vocabulary that has become the “gold standard” for annotation of the small molecular entities. |
| **CHEMINF** | CHEMical INformation ontology<sup>48</sup><br>Can be used to annotate chemical information, including chemical structure and chemical properties. Importantly, it includes many of the database and structure identifiers such ChEMBL ID or InChIKey, respectively. |
| **OBI** | Ontology for Biomedical Investigations<sup>49</sup><br>Can be used to describe a biomedical investigation, such as protocols, instruments and experimental design. Importantly, it allows to annotate data points of specific doses used in the experiment (http://purl.obolibrary.org/obo/OBI_0000984) that examine dose-response (http://purl.obolibrary.org/obo/OBI_0001418) relationships. Furthermore, it can be used explicitly to annotate an *in vitro* (http://purl.obolibrary.org/obo/OBI_0001285) experiment. |
| **CLO** | Cell-line Ontology<sup>50</sup><br>CLO is useful for describing a cell-line. It aims to bring together available cell-line data from multiple sources into a common format<sup>50</sup>. Importantly, it can be used to specify a cell-line that was maintained in an *in vitro* environment (http://purl.obolibrary.org/obo/CL_0001034), thus allowing identification of *in vitro* investigation. |
| **Cellosaurus** | The Cellosaurus: a cell line knowledge resource<sup>33</sup><br>Cellosaurus is used for referencing cell-lines. As of version 24 (November 2017) it contained 99 021 cell-lines with references to original publications and cross-references to many cell-line catalogues. They also provide references to resources, where samples (BioSamples, ChEMBL, etc.) could be annotated with the cell-lines contained in Cellosaurus. Similarly, it allows search of *in vitro* studies through the identification of specific cell-lines. |
| **BAO** | BioAssay Ontology<sup>51</sup><br>BAO was developed to describe and model diverse and complex assays <sup>51</sup>. Importantly, BAO provides definitions for absolute or relative IC50 (http://www.bioassayontology.org/bao#BAO_0000197, http://www.bioassayontology.org/bao#BAO_0000198, respectively) values and for several other endpoints based on bioassay measurements. |
| **UO** | Units of Measurement<sup>52</sup><br>UO provides units of measurement. Importantly one can use units of molarity (http://purl.obolibrary.org/obo/UO_0000061) such as micromolar (http://purl.obolibrary.org/obo/UO_0000064) to define concentrations in *in vitro* experiments, thus allowing identification of *in vitro* data. |
| **UBERON** | Uberon, an integrative multi-species anatomy ontology<sup>53</sup><br>UBERON is an integrated cross-species ontology that facilitates reasoning from higher-order anatomical levels to more specific sublevels. For example, CLO uses UBERON to add organ-specific annotations to cell-lines<sup>50</sup>. Therefore, UBERON can be used to translate knowledge from cell-lines to organs or vice versa. |
| **EDAM** | EDAM: an ontology of bioinformatics operations, types of data and identifiers, topics and formats<sup>54</sup> |
|  | EDAM can be used to describe an *in vitro* scientific experiment, such as the output and computational procedures related to the data and data formats. Similar to CHEMINF it provides vocabulary for compound structure identifiers/format such as InChIKey (http://edamontology.org/format_1199), however it is more useful in annotating the down-stream data analysis and less practical in describing the experimental design. |
| **EFO** | Experimental Factor Ontology<sup>55</sup> |
|  | EFO was developed to annotate the large quantity of domain-independent gene expression data<sup>55</sup> stored in ArrayExpress. Although not specifically developed for use with chemical compounds, it contains specific classes describing chemical entities (e.g. chemical compounds and drugs). EFO ontology can be used to identify gene expression data that is related to chemical compounds. |

### Using ontologies to retrieve *in vitro* compound data

Since the information stored in an RDF triplet is formulated as subject-predicate-object, each element can be a term defined in an existing ontology. Therefore, we investigated if using ontologies enabled us to retrieve samples directly related to *in vitro* ontology terms. We used BioSamples public SPARQL endpoint as our target database since it combines sample annotations from ArrayExpress, PRIDE Archive and European Nucleotide Archive (ENA) and as shown previously, we were able to retrieve data from BioSamples for all our model compounds using both compound and IUPAC systematic names. To our surprise, none of the samples in BioSamples were annotated with the *in vitro* ontology term (http://purl.obolibrary.org/obo/OBI_0001285) found in Ontology for Biomedical Investigation (OBI) ontology or with structural identifier ontology terms (InChI, InChIKey, SMILES) from either ChEBI (Chemical Entities of Biological Interest), CHEMINF or EDAM (Bioinformatics operations, data types, formats, identifiers and topics) ontologies. However, we did find samples for all our model compounds using ChEBI ontology terms (rosiglitazone: http://purl.obolibrary.org/obo/CHEBI_50122, aspirin: http://purl.obolibrary.org/obo/CHEBI_15365, valproic acid: http://purl.obolibrary.org/obo/CHEBI_39867, tamoxifen: http://purl.obolibrary.org/obo/CHEBI_41774).

Next, we investigated if samples of our model compounds retrieved using ChEBI ontology terms were associated with a molarity unit (http://purl.obolibrary.org/obo/UO_0000061) using the UO (Units of measurement) ontology. Indeed, such samples existed for *rosiglitazone* and *tamoxifen* but not for *aspirin* and *valproic acid*. Alternatively, we were interested if any of the samples associated with the model compounds were also associated with a cell-line ontology term (http://www.ebi.ac.uk/efo/EFO_0000322) from Experimental Factor Ontology (EFO). No samples were found for *aspirin* but we were able to find several samples for *rosiglitazone* and *tamoxifen* and a single sample for *valproic acid* (Supplementary Tables 10-12), effectively identifying *in vitro* compound data.

In summary, despite the fact that we were able to retrieve some samples representing *in vitro* experiments related to our model compounds, they are of limited use for the search of presently available compound data, as these data are mostly deposited without associated ontology terms. However, the use of ontologies could be a powerful tool for search strategies once annotations of experimental metadata improves.

## Discussion

In this study, we used four model compounds (*aspirin, rosiglitazone*, *valproic acid* and *tamoxifen)* to investigate 19 public resources potentially suited for identification and access to *in vitro* compound data. Fragmentation and problematic accessibility of *in vitro* data poses a major obstacle for their optimal use and hence reduces their scientific and practical impact. Quick and precise identification and information retrieval is necessary for efficient use of published data, to save researcher and regulators alike valuable time and to allow quick and easy identification of relevant datasets^56^.

## Fragmentation of public resources and integration efforts

There are many resources that provide compound-specific data. There are also several databases that contain omics data. A general consensus is that raw data from experiments should be publicly accessible, and it is the policy of high quality scientific journals and of several granting agencies to require publicly accessible deposition of research data. In theory, this is an excellent opportunity for both *in vitro* and *in silico* toxicology since it would allow to compare data produced by different research laboratories and help to estimate the impact of different experimental conditions. However, at present, this theoretical opportunity does not live up to its full potential.

Efforts have been made to integrate compound information or omics data from several resources. Indeed, in our analysis we observed that many curated resources dedicated to chemical compounds contain cross-references to each other. Some of them also include data from one another. This can lead to databases overlapping in their content^57^. As an example of cross-references, ChemSpider reports to have links to more than 500 resources but none representing an omics data resource. On the other side, BioSamples database integrates metadata from omics resources such as ArrayExpress, European Nucleotide Archive (ENA) and PRIDE but does not readily associate this metadata with compound-specific resources or compound identifiers. Although resources such as UniProt, ChEMBL and ChEBI included links to ArrayExpress, ExpressionAtlas or PRIDE, we observed that these were not exhaustive, with several potential links to omics datasets missing.

In addition to BioSamples, there are other public and private initiatives aiming at bridging the gap between the discrepancies of field-specific databases. OmicsDI (Omics Discovery Index)^58^ and Repositive IO (https://repositive.io) aim to integrate metadata from many different domains, including genomics, transcriptomics and metabolomics. Although they lack cross-references to compound-specific resources, they can still be used to identify compound-related *in vitro* omics data. Furthermore, Repositive IO allows its community driven search platform to be populated with annotation tags. This makes data identification a crowd-sourcing task, which may substantially improve the annotation of the metadata. However, currently these annotations suffer from the lack of a common vocabulary and may lead to confusion when different terms that correspond to the same meaning are used. Instead, a system using standardized ontologies should be favored. In that case compound samples could be directly related to the compound’s InChIKeys as an annotation tag describing the metadata. Similar additions could be made with concentration units and cell-lines.

## Variability of compound names and identifiers

It is well known that free-text search of systematic chemical names has low precision and high error ratei^12^, especially considering the variability in synonyms between resources as demonstrated for our model compounds. Therefore, searching a compound name or its synonym from one resource might fail to retrieve results from another resource. Also, covering the whole scope of compound identifiers will take substantial amount of researcher’s time, since a single compound can contain hundreds of synonyms and several identifiers. Furthermore, compound names themselves are ambiguous, especially in omics resources that are not specifically designed to store compound information where associated metadata fails to incorporate unique compound identifiers. For example, as identified in our study, *measurin*, which according to ChEMBL is a synonym of *aspirin* (Compound ID: CHEMBL25), can potentially be misled with samples where labels include the term *measuring*. Indeed, two out of four model compounds in our study were annotated using alternative labels that were associated with different samples.

Ideally, only a single unique structure representation exists for each unique compound. We tested this by using the name of the model compound as search input and recorded four structural identifiers: InChI, InChIKey, SMILES and IUPAC name. Most variability was observed for SMILES and IUPAC names where more complex molecule, such as *rosiglitazone* and *tamoxifen* had multiple structural identifier representations between resources. In the case of InChI and InChIKey, the variation, was considerably lower, albeit not absent. Therefore care must be taken not to attribute bioassay results to alternative stereochemical structures^57^.

The observed variability in structural identifiers is not surprising, since often molecular formulas are not unequivocally unique representations of a chemical compound^39^. Nevertheless, although several InChIKeys can be derived from a single InChI, they are still considered sufficiently unique at providing an adequate collision resistance i.e. it is unlikely that two different compounds are associated with one InChIKey^59^. However, including all omics resources, there are some compound-specific resources that don’t allow data identification based on compound’s structural identifiers. This can cause problems for data integration tasks which is why compounds should always be supplemented with unique structural identifiers, such as InChIKeys, that would enhance their downstream identification.

## Using ontologies to integrate and search for *in vitro* compound data

One way to increase annotation quality is to use existing ontologies that enhance the quality and consequently the precision of identifying data correctly. In addition, since ontologies are a part of the “linked data” concept, they can be used to integrate information from several resources by using federated queries. To this end, ontology browsers, such as Ontology Lookup Service (OLS)^36^ are excellent tools for finding terms relevant for your data, thus also promoting the reuse of existing ontology content^35^.

In this study, we focused on resources that provided Resource Description Framework (RDF) compliant data or a public SPARQL endpoint for data queries since they inherently take advantage of ontologies. We failed to retrieve a single sample in BioSamples resource that would directly relate to *in vitro design* ontology term from Ontology for Biomedical Investigation (OBI) ontology. In addition, we searched BioSamples for datasets characterized with molar concentration unit or cell-line ontology terms. Similarly, the results were far from comprehensive. We think that this approach can be further improved once better ontology mappings of the metadata become available. Prospectively, there is a clear potential, provided a better annotation of data, for example by utilizing the Cellosaurus ontology ^33^ that contains a controlled vocabulary for around 100 000 cell lines.

Using SPARQL and other semantic web technologies, we showed that associations between raw data samples and ontology terms do exist and hence the ambiguity of identifying and retrieving *in vitro* data, can be reduced. Nevertheless, despite the current state of data annotation, in order to integrate data from different public resources, the compound formatting needs to be further harmonized and the domain vocabulary standardized^56^. More importantly, the new and existing resources need to adjust themselves to take full advantage of existing standardized ontologies. So far, they have only been implemented by the selected few.

To a certain extent, there are initiatives that utilize semantic web technologies to solve problems with compound data integration. As such, the Open PHACTS project, encourages data resources to publish their data as RDF, in order to build an “open pharmaceutical space” where data can be accessed through a user-friendly software interface^10^. As of today, it is not clear how the software can be used by researchers who do not have explicit knowledge in API programming. Nevertheless, it can potentially be used to build new, researcher friendly applications on Open PHACTS interoperable data. In parallel, OpenTox aims to provide a single access to both *in vivo* and in *vitro* toxicity data and a framework for storing and executing predictive computational models^60^. Similar to Open PHACTS, this initiative is not applicable to non-expert researcher, who is looking for compound associated raw omics data, thus a gap connecting compounds to their raw *in vitro* data still persists.

## *In vitro* toxicology: the magnitude of the challenge ahead (conclusion and out-look)

Tens of thousands of compounds need to be tested in order to correctly evaluate the pharmaceutical chemical space^61^. Realizing this task using *in vivo* models is expensive and time consuming. Instead, alternative test methods such as high-throughput *in vitro* assays provide the capability to test thousands of compounds across a wide range of concentrations. Together with predictive *in silico* applications, such as Toxtree^62^, which is an open-source application commissioned by the European Commission Joint Research Center’s European Chemical Bureau, they have the potential to improve risk assessment of chemicals for the benefit of human health and environment.

There already exists a substantial corpus of resources that contain data on a large number of chemical compounds. These data and their sources are diverse and they need to be integrated in order to answer the systemic effects of compounds (Figure 2). Accessing published data with correct compound information is essential. The problems encountered in accessing data on our model compounds, demonstrate that using the results from publications stored in public resources and cross-referencing them with omcis data still requires substantial investigative capacity. We recognize, that there are two solutions for the problems of accessing *in vitro* activity data: curation of already existing data and improved annotations by the experimenters. Curation could transform poorly and hard to reach compound information into a rich and relevant resource. Without new experimental efforts, it could right away enhance our knowledge of toxicology and decrease unnecessary additional experimentation, including animal testing. The problem of the curation is unfortunately in large parts insurmountable: the manpower and expertise required to transform the present “data graveyard” into a living resource would necessitate financial means which simply do not exist. Thus, we fully support the concept of a curation effort but we are also perfectly aware of its limitations. However, efforts similar to SourceData^63^, that allows to annotated already published figures in publication, could provide a potential cure. Alternative to curation is improved annotation by the experimenter themselves. Obviously, this is a prospective effort that will not apply to existing data. It is also limited by the willingness and expertise of the experimenter to standardize data presentation. For example, the use of proper compound identifiers (e.g. InChIKeys) and ontology terms is not part of the present scientific culture. However, the use of ontology browsers could alleviate this transition. Nevertheless, currently researchers receive little to no reward for the broad accessibility and perennity of their data. Thus, while improved annotation by the experimenter is a noble goal for the future, this will not happen automatically. Novel approaches for scientific publishing such as the use of proper annotations with ontology terms and attribution of credit to scientists for their data will be needed to really make an impact. Would the efforts necessary for general accession to *in vitro* compound data be worth the money and time? Considering the success of UniProt which incorporates extensively curated and trustworthy protein data, the answer is yes. Indeed a careful analysis published in the EMBL-EBI value report^64^ estimated 46% increase in research efficiency for scientists accessing information relevant to their research question. With around 400 000 unique visitors per month, the reported estimation shows an enormous cost-effect benefit for the researcher community. The interest in chemical compounds is even bigger: PubChem alone receives about 1 million unique users per month^65^. However, omics resources do not readily link to PubChem (or other compound databases) and often ambiguous compound names are used to annotate the data. Thus, the time lost by the community to assemble compound information is considerable. This highlights the need for an improved resource that would enhance the efficiency and speed of accessing raw and analyzed compound data in a reliable, simplified and intuitive manner. It would allow researchers to focus on data analysis and its interpretation instead of collection and curation. And finally, such a resource could contribute to public health by allowing a better identification and management of potentially hazardous compounds.

**Figure 2.**
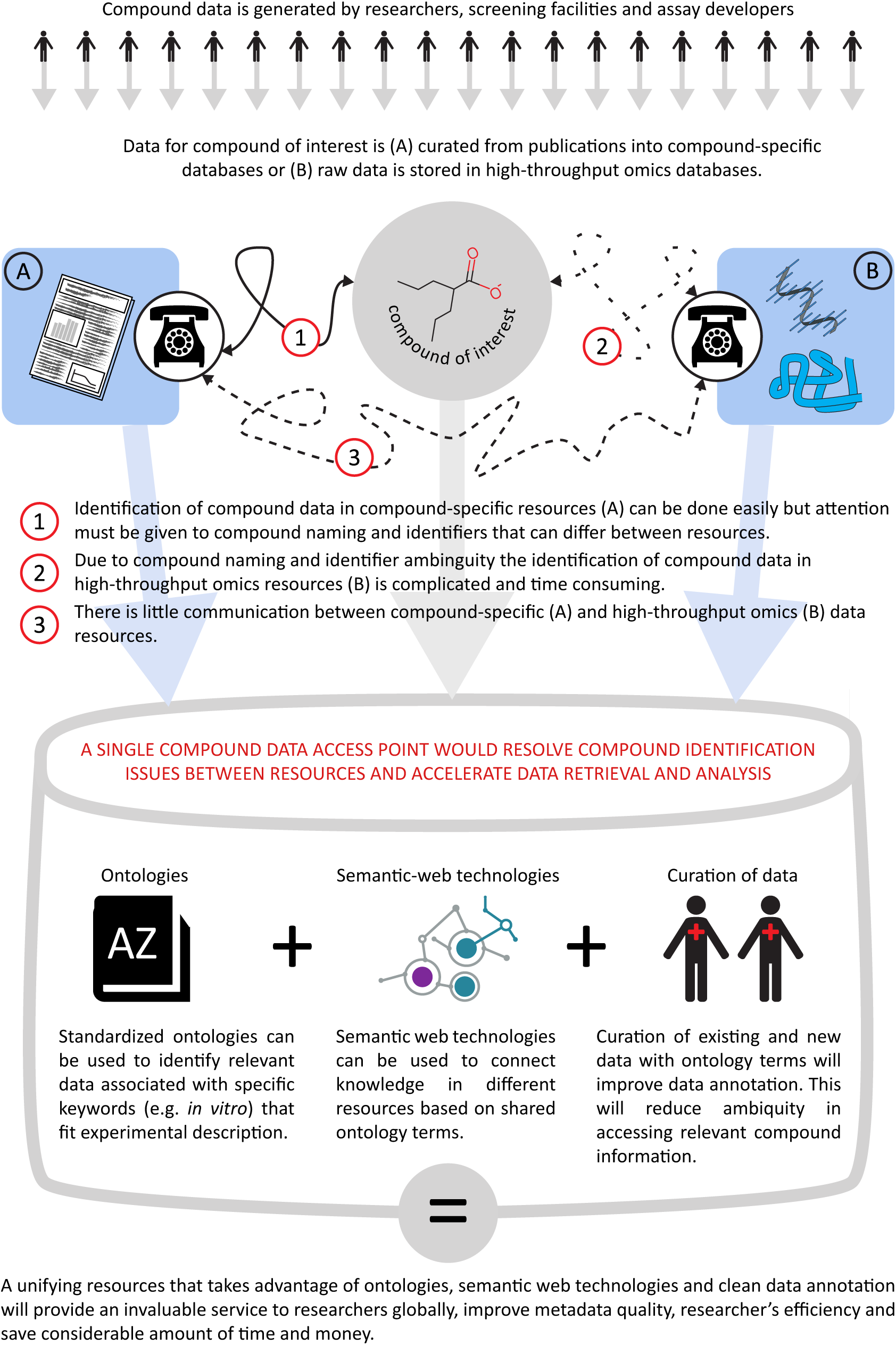
A graphics illustrating the problems of integrating knowledge between compound of interest and different types of data resources. The problems can be solved with integrated approaches using ontologies, semantic-web technologies and better annotation of the data.

## Acknowledgements

SI and JS were partially supported by Swiss Centre of Applied Human Toxicology SCAHT).

